# Phylogenetic distribution of secondary metabolites in the *Bacillus subtilis* species complex

**DOI:** 10.1101/2020.10.28.358507

**Authors:** Kat Steinke, Omkar S. Mohite, Tilmann Weber, Ákos T. Kovács

## Abstract

Microbes produce a plethora of secondary metabolites that although not essential for primary metabolism benefit them to survive in the environment, communicate, and influence differentiation. Biosynthetic gene clusters (BGCs) responsible for the production of these secondary metabolites are readily identifiable on the genome sequence of bacteria. Understanding the phylogeny and distribution of BGCs helps us to predict natural product synthesis ability of new isolates. Here, we examined the inter- and intraspecies patterns of absence/presence for all BGCs identified with antiSMASH 5.0 in 310 genomes from the *B. subtilis* group and assigned them to defined gene cluster families (GCFs). This allowed us to establish patterns in distribution for both known and unknown products. Further, we analyzed variations in the BGC structure of particular families encoding for natural products such as plipastatin, fengycin, iturin, mycosubtilin and bacillomycin. Our detailed analysis revealed multiple GCFs that are species or clade specific and few others that are scattered within or between species, which will guide exploration of the chemodiversity within the *B. subtilis* group. Uniquely, we discovered that partial deletion of BGCs and frameshift mutations in selected biosynthetic genes are conserved within phylogenetically related isolates, although isolated from around the globe. Our results highlight the importance of detailed analysis of BGCs and the remarkable phylogenetically conserved errodation of secondary metabolite biosynthetic potential in the *B. subtilis* group.

**IMPORTANCE:** Members of the *B. subtilis* species complex are commonly recognized producers of secondary metabolites, among those the production of antifungals makes them promising biocontrol strains. However, while there are studies examining the distribution of well-known *B. subtilis* metabolites, this has not yet been systematically reported for the group. Here, we report the complete biosynthetic potential within the *Bacillus subtilis* group species to explore the distribution of the biosynthetic gene clusters and to provide an exhaustive phylogenetic conservation of secondary metabolite production supporting the chemodiversity of *Bacilli*. We identify that certain gene clusters acquired deletions of genes and particular frame-shift mutations rendering them inactive for secondary metabolite biosynthesis, a conserved genetic trait within phylogenetically conserved clades of certain species. The overview presented will superbly guide assigning the secondary metabolite production potential of newly isolated strains based on genome sequence and phylogenetic relatedness.

## OBSERVATION

*Bacilli* can be isolated from various environments, plant rhizosphere, animal and human digestive system, where secondary metabolites (SMs), metabolites not necessary for primary metabolism, play a pivotal role. The *Bacillus subtilis* group, which includes *B. subtilis* and its closely related species (Fig. 1), comprises common producers of bioactive SMs such as antimicrobials and cytotoxic substances, empowering them for a range of industrial applications, including plant pathogen biocontrol (1, 2). Members of the *B. subtilis* group are producers of numerous well-known natural products, iturin, mycosubtilin, fengycin/plipastatin, or bacillaene. Only recent studies have emerged that investigated species-level distribution of the corresponding biosynthetic gene clusters (BGCs) in the *B. subtilis* group (3). Recent reviews provide an overview of various SMs produced by these *Bacilli (1, 4)*.

**Fig. 1:**
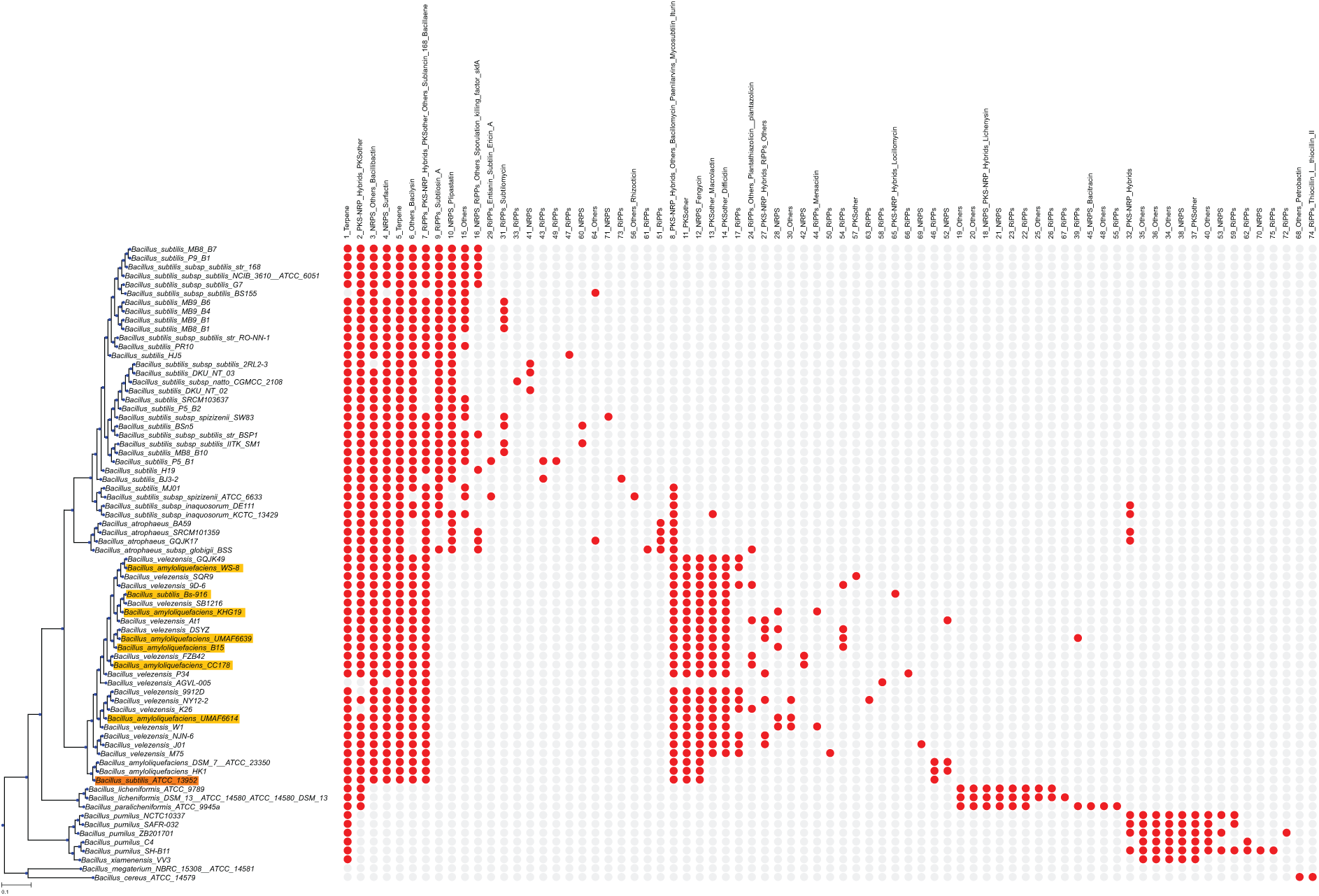
Phylogenetic tree based on a multilocus sequence alignment of 30 genes with a modified version of autoMLST, using IQ-TREE and ultrafast bootstrapping with 1000 replicates. *B. cereus* ATCC 14579 and *B. megaterium* NBRC 15308 were used as an outgroup. Absence/presence of GCFs is visualized with a red dot indicating presence and a gray dot indicating absence. Fig. S1 includes the complete tree. Strains with disagreements in NCBI and GTDB taxonomy are highlighted.

Here, we significantly expand previous studies by investigating patterns in all complete *B. subtilis* group genomes as of July 2019. We therefore examined phylogenetic distribution of BGC families in 310 *B. subtilis* group genomes (Data S1) by predicting BGCs with a modified version of antiSMASH 5 (5), clustering these into gene cluster families (GCFs) with BiG-SCAPE (6), and visualizing GCF distributions in a phylogenetic tree generated with autoMLST-derived scripts (7) (Fig. S1, S2).

Phylogeny was generated based on a multilocus sequence alignment of 30 conserved single-copy genes (Fig. 1), generally reflecting NCBI taxonomy, but with certain disagreements in the *B. velezensis* and *B. amyloliquefaciens* clades (highlighted in Fig. 1 and Materials and Methods).

The 3655 BGCs identified using antiSMASH 5 (5) were assigned into 75 GCFs and 62 singletons with BiG-SCAPE (6) (Fig. S2), GCFs were subsequently mapped to the tree (Fig. 1). Only one predicted metabolite, a terpene (sesquarterpene), was found in nearly all strains, while another, a predicted NRPS/PKS hybrid, was found in most species except *B. pumilus* and *B. xiamenensis*. Other widespread GCFs are bacillibactin, surfactin, and bacilysin. Bacillaene and sublancin 168 families were found in most species, except *B. licheniformis, B. paralicheniformis, B. pumilus*, and *B. xiamenensis*; however, there are two gaps seemingly following clade boundaries in *B. subtilis*. A similar gap in distribution occurs in bacilysin, which is absent in *B. spizizenii* and *B. atrophaeus*.

Such apparently clade-linked patterns in absence or presence of GCFs were common - many GCFs were distributed according to phylogeny, either linked to clades spanning multiple species or limited to a single species. Lichenysin, a clade-specific GCF, was identified only in *B. licheniformis* and *B. paralicheniformis*. Distribution of the highly similar lipopeptides fengycin and plipastatin also followed clade boundaries, with fengycin in *B. velezensis* and *B. amyloliquefaciens* and plipastatin in *B. subtilis* and *B. atrophaeus*. However, as previously reported (8), no plipastatin was found in *B. spizizenii*, but rhizocticin, supporting clade’s biosynthetic distinctness.

Other examples of clusters almost or entirely limited to one species in the tree included bacitracin, which was present in all examined *B. paralicheniformis* genomes, and difficidin and macrolactin, both found in most *B. velezensis* (though macrolactin was also present in single isolates of other species). Certain species-specific GCFs were found dispersedly, for instance, the *B. subtilis*-specific subtilomycin was apparently linked to certain clades within the species, or the ribosomally synthesized and post-translationally modified peptide coding 33_RiPP or 49_RiPP families that were also species specifics. Additionally, certain families appeared in multiple clades, but in a clade-linked pattern (17_RiPPs in *B. velezensis*), while others were missing in one or more clades (15_Others in *B. subtilis*).

Finally, other GCFs appeared more scattered within a species, with no link to a certain clade evident, such as the 42_NRPS GCF in *B. velezensis* (Figure S1). Only a few GCFs (e.g. 39_RiPP), appeared scattered across the entire tree without noticeable link to certain clades. Scattered patterns and random occurrences of GCFs outside of key species might be driven by horizontal gene transfer, in accordance with natural competence of *B. subtilis* (4) for instance, the 42_NRPS family contains the *nrs* cluster of *B. velezensis* FZB42, suggested to be acquired via HGT (9).

Next, we compared the genetic variation within particular GCFs and investigated the phylogenetic relation among these variants, selecting families that code for important *Bacillus* SMs, fengycin, plipastatin, iturin, bacillomycin, and mycosubtilin.

A total of 128 BGCs were part of the similarity network with BGCs for plipastatin, a biodegradable fungicide (1). Based on the similarity network these BGCs were grouped into 6 groups: PPS, PPS groups B to E and PPS_others (Fig. 2, Data S2). Plipastatins are mostly observed in the *B. subtilis* genus, with exception of group B BGCs in *B. atrophaeus*. We found that 72 BGCs from group PPS and 7 BGCs from group B had all biosynthetic genes (BGs) for plipastatin (*ppsA* to *ppsE*). In contrast, groups C, D, E and “others” had BGCs missing upto three BGs (Fig. S3), consistent with experimental data demonstrating lack of plipastatin production in *B. subtilis* natto BEST195 (10) and *B. subtilis* P5_B2 (3) (Fig. 2). A similar deletion of BGs was found in several other strains. Interestingly, these strains are phylogenetically close to each other suggesting such deletions being conserved within a single clade (Fig. 2F, Fig. S4). Additionally, in plipastatin BGCs of group E, gene *ppsE* appeared to have missing domains (Fig. S5). Investigation of the nucleotide sequences of *ppsE* gene homologs revealed a deletion at position 232 of reference *ppsE* gene across all the 16 members of group E, leading to frameshift (Fig. S5). This frameshift was present in multiple strains isolated from distincts geographic locations (Data S2) but belonging to the same phylogenetic clade suggesting an evolutionarily conserved frameshift in *ppsE* gene that could lead to loss of function.

**Fig. 2.**
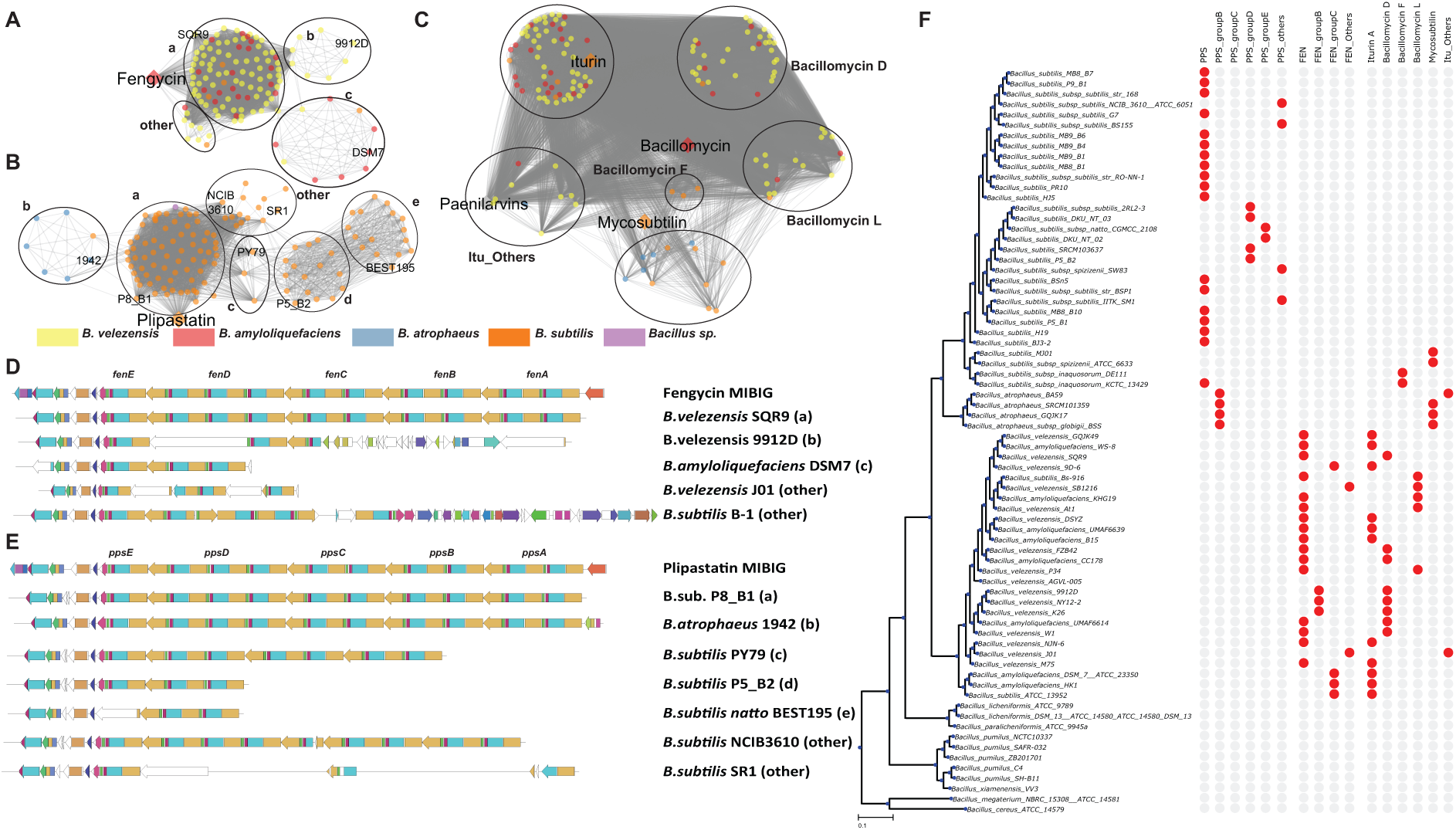
Comparison of plipastatin/fengycin/iturin families of BGCs. A-C. Similarity networks representing the plipastatins, fengycins, and iturin-like BGC families, respectively. The different colors represent different species of *Bacillus*. Iturin like BGCs are grouped based on amino acid specificity predictions instead of BiG-SCAPE generated similarity index (Table S1). D-E. Selected clusters are shown from different groups of plipastatins and fengycins, respectively. The detailed genetic structure of all incomplete BGC families can be found in Fig. S3. F. Phylogenetic distribution of different groups of BGCs is presented across selected genomes. For a complete list of all genomes and different groups of BGCs, see Fig. S4.

The fengycin family network contained 123 BGCs from *B. velezensis* and *B. amyloliquefaciens* species, in addition to 5 isolates likely misclassified as “*B. subtilis”*, which based on MLSA data should be assigned as *B. velezensis* strains (Data S2, Fig. S4). The fengycin BGCs could be divided into 4 groups, with 96 BGCs containing all BGs (*fenA* to *fenE*). BGCs from groups B, C and others contained incomplete BGCs with upto three of the BGs missing (Fig. S3). The strains harboring these incomplete fengycin BGCs were also found to be phylogenetically close, similar to plipastatins, suggesting these deletions being conserved within a single clade (Fig. 2F, Fig. S4). As above for *ppsE* gene, many phylogenetically close strains harboring group B of fengycin contained a frameshift at positions 3126-3127 of *fenD* gene suggesting a possible evolutionary trait of the clade (Fig. S6).

Unlike above, sequence similarity alone could not divide the 141 iturin-like BGCs into distinct groups due to conserved BG sequences, which differ only in the individual amino acid substrate specificities leading to production of diverse lipopeptides like iturin A, bacillomycin D-F-L, and mycosubtilin (11) different levels of bioactivity (4). Therefore, antiSMASH predicted amino acid substrate specificity for all NRPS adenylation domains was used to group the BGCs into iturin A, bacillomycin D-F-L, mycosubtilin and “others” that have less than 7 (typical of iturins) amino acid substrates (Table S1, Data S2). Mapping these data onto the phylogenetic tree revealed that each group is conserved in closely related strains. Mycosubtilin group detected in *B. atrophaeus* and few *B. subtilis* (mostly in subspecies *spizizenii*), bacillomycin in 3 of *B. inaquosorum*, whereas iturin A, bacillomycin D and bacillomycin F was spread across different *B. velezensis* and *B. amyloliquefaciens* isolates, confirming previously proposed species and strain level presence of iturinic lipopeptides (11).

Our detailed BGC comparison identified variations in particular GCFs to be phylogenetically conserved. Such phylogenetic correlation of different BGC groups and particular frameshifts suggest evolutionary relationships among production capabilities of *Bacillus* strains. Therefore, our workflow combining comparative analysis of BGCs and phylogenetic relationships revealed how a particular BGC is evolved within species. This knowledge, and closer examination of the exceptions, may guide selection of specific strains as antimicrobial producers within underexplored groups of SM producers.

## MATERIALS AND METHODS

### Genome selection

Initially, all genomes of *B. amyloliquefaciens, B. atrophaeus, B. licheniformis, B. paralicheniformis, B. pumilus, B. subtilis, B. velezensis, B. xiamenensis* and a few related *Bacillus* sp. strains with assembly status “complete” or “chromosome” publicly available from NCBI in July 2019 were selected. Additionally, the type strains of *B. cereus* and *B. megaterium* were included as outgroups. The strain list was then curated to remove duplicates. Further, the genomes of engineered *B. subtilis* and strains were removed(*B. subtilis* BEST7613, *B. subtilis* delta6, *B. subtilis* IIG-Bs27–47-24, *B. subtilis* PS38, and *B. subtilis* PG10, as described in (12), as well as *B. subtilis* BEST7003, *B. subtilis* QB5413, *B. subtilis* QB5412, *B. subtilis* QB928, and *B. subtilis* WB800N). After preliminary tree reconstruction, *B. subtilis* HDZK-BYSB7 was found to clade with *B. cereus* rather than the other *B. subtilis* strains and was therefore removed; it has since been reclassified as *B. anthracis*. Initial examination of results also found BGCs to be split by the origin in *B. velezensis* Hx05; for ease of analysis, this strain was therefore dropped. Subsequently, *B. velezensis* AGVL-005 was found to contain many frame-shifted proteins; however, it was retained. A further 13 in-house genomes of *B. subtilis* and one of *B. licheniformis (13)* were included. This led to a final count of 310 genomes.

### Genome acquisition and strain name annotation

Genomes were downloaded in genbank format with the ncbi-acc-download tool (https://github.com/kblin/ncbi-acc-download). As many of the genbank entries did not contain strain information in the “Source” or “Organism” features, which are required by the autoMLST and BiG-SCAPE tools to distinguish the individual strains, the Python script rename_strainless_organisms.py (found in the tree and matrix construction pipeline, see below) was employed to transfer strain information from the “strain” field to these fields.

### Genome mining

In a first step, all downloaded genomes were initially mined for SMs with antiSMASH 5.0 (5). As antiSMASH collapses gene clusters that are encoded in close proximity, such as the iturin and fengycin clusters in *Bacilli*,into a single biosynthetic “region”, a modified version of antiSMASH (https://github.com/KatSteinke/dmz-antismash) was developed that contains the additional functionality to split known clusters at a user-defined gene, resulting in two independent “regions”. In all other respects, this version of antiSMASH is identical to antiSMASH 5.0.0. The modified version of antiSMASH was run as an antiSMASH fast run with the default parameters. The genes selected to split between adjacent clusters were *dacC* and yngH for the plipastatin/fengycin clusters, and *yxjF* and *xynD* for the iturin clusters. For assigning the plipastatin/fengycin boundary genes, homologs from several species were selected to reflect species variations: *dacC* homologs from *B. velezensis, B. subtilis, B. amyloliquefaciens* and *B. atrophaeus* and *yngH* homologs from *B. subtilis* and *B. atrophaeus*. These were selected so that the cut would yield the intersection of both clusters as found on MIBiG, from *dacC* to *yngH*, as other boundaries led to incorrect splits, either failing to cut the cluster or cutting it twice. The genes are identified by a BLAST search in the examined genome, with coverage and identity of at least 90% each needed for identification. During this step, errors in the Genbank file of *B. licheniformis* PB3 (NZ_CP025226.1) were detected, as they caused subsequent errors in antiSMASH; the erroneous portions, CXG95_RS00005 and CXG95_RS00010, were consequently deleted.

### GCF identification and clustering

To identify families of homologous gene clusters present in multiple species (gene cluster families, GCFs), BiG-SCAPE (6) was used at default settings. In order to automatically identify any known compounds, reference clusters from the MIBiG database (14) were included in the networking analysis. Singleton clusters were not returned. As this produced almost exclusively GCFs split along species lines, even for compounds known to be found in all species, connected components were identified with the NetworkX library (15) using a similar approach as in (16). However, as BiG-SCAPE was left at default options, duplicated entries were later merged.

### Tree building

For getting a highly resolved phylogeny of the closely related *Bacillus* strains, maximum likelihood trees were constructed with a pipeline based on autoMLST (7), and using autoMLST defaults to the greatest extent. A modification was introduced to autoMLST that skipped the automated search and inclusion of similar genomes and thus only processed the supplied genomes. Subsequently, the pipeline identifies all conserved single-copy genes from these genomes. Additionally, the gbk2sqldb.py script in autoMLST, which was employed in the pipeline, was patched to use the same hmm database (reducedcore.hmm) as the main automlst.py script. The modified version is available at https://github.com/KatSteinke/automlst-simplified-wrapper.

Both for the short tree shown in Fig. 1 and the full tree (Fig. S1), this yielded 30 single-copy/housekeeping genes for each tree; however, not all of these were identical between the trees. For generating the multi-locus alignment, each individual gene was aligned with MAFFT (17) and the alignment trimmed using trimAl (18); then, all alignments were concatenated. As in autoMLST, the tree was generated with IQ-TREE (19), using Ultrafast Bootstrap (20) with 1000 replicates.

The resulting tree was rerooted in ETE3 (21) during the visualization step, using *B. megaterium* NBRC 15308 and *B. cereus* ATCC 14579 as an outgroup.

Based on our analysis, in line with the recently released genome-based taxonomy in GTDB (22, 23), these strains should be designated *B. velezensis* or *B. amyloliquefaciens*, respectively. In our global analysis of all 310 genomes, a total of 28 strains whose genome-based taxonomy conflicts with their assigned species names were identified (Fig. S1, Data S1). Additionally, strains designated *B. subtilis* subsp. *inaquosorum* and *B. subtilis* subsp. *spizizenii* by NCBI form their own clades, consistent with their recent promotion to species status. (8) The tree thus appears to reflect genome-based taxonomy well.

### Absence/presence matrix

An automated tree and matrix construction pipeline was established that tied together the individual steps of the analysis. The script for this pipeline takes as arguments the location of a base directory in which analysis results are to be placed, the location of a file listing accession numbers to be downloaded, the name of the final tree to be output and optionally outgroups to be used. It creates all the files and directories necessary for the subsequent analysis (see below). The script can be downloaded at https://github.com/KatSteinke/AbsPresTree.

From the connected component GCFs, a matrix counting occurrence of each GCF in each strain was computed. GCFs were subsequently clustered according to their occurrence in each strain using SciPy’s clustering package (24); hierarchical clustering was performed. Subsequently, the absence/presence matrix was reordered to reflect the clustering of GCFs. It must be noted, however, that the connected component GCFs are based on placement of gene clusters in a network, and even incomplete or inactive clusters may be included if they pass the threshold for clustering. The tree and matrix were visualized in ETE3 using ETE3’s clustering module. Subsequently, matrix columns were manually arranged to follow phylogeny of the strains primarily represented per column.

### Variations within particular GCF

Based on the similarity networks of plipastatin and fengycin GCF, we created groups within a GCF. The fengycin GCF was split into 4 groups and plipastatin GCF into 6 groups. The genetic structure variations among groups with few missing BGs are shown in Figure S3. The genes *ppsE* from plipastatin group E and *fenD* from fengycin group B are further selected for multiple sequence alignment (Fig. S5 and S6). For the iturin-like lipopeptide GCF, substrate specificities of A-domain were collected from antiSMASH annotations. Based on the individual amino acid specificities the BGCs from this GCF are further classified into iturin A, bacillomycin D, F, L and mycosubtilin (Table S1). The script used to analyze the variations in GCF can be downloaded at https://github.com/OmkarSaMo/GCF_variation_Bacillus.

### Data availability

The data with NCBI accession IDs and information on all detected gene clusters is available in supplementary Data S1 and S2. Code used to generate the data is available at https://github.com/KatSteinke/AbsPresTree. The script used to analyze the variations in GCF is available at https://github.com/OmkarSaMo/GCF_variation_Bacillus.

## ACKNOWLEDGMENTS

This work was funded by the Danish National Research Foundation (DNRF137) for the Center for Microbial Secondary Metabolites. T.W. and O.S.M. furthermore acknowledge funding from the Novo Nordisk Foundation (NNF10CC1016517, NNF16OC0021746).

## AUTHOR CONTRIBUTIONS

KS and OSM performed the bioinformatic analysis; KS, OSM, TW, and ATK interpreted the data; KS, OSM, TW, and ATK wrote the manuscript.

## SUPPLEMENTAL MATERIAL

**Fig S1** Phylogenetic tree and presence absence of different BGC families across *Bacillus* group

**Fig S2** Similarity network overview of all BGCs detected across *Bacillus* genome visualized using Cytoscape (Shannon P, Markiel A, Ozier O, Baliga NS, Wang JT, Ramage D, Amin N, Schwikowski B, Ideker T, Genome Res 13:2498–2504, 2003).

**Fig S3** Genetic structure variations in partial BGCs of plipastatin and fengycin families

**Fig S4** Phylogenetic tree distribution of different fengycins and iturins across 310 genomes of *Bacillus sp*.

**Fig S5** Conserved frameshift in the *ppsE* gene across strains harboring group E plipastatin BGCs. A. BGCs from group E of plipastatins with missing domains in *ppsE* gene. B. Part of the phylogenetic tree including plipastatin group E strains (Complete tree in Figure S4). C. Nucleotide sequence alignment with conserved deletion at position 232 that lead to frameshift. The nucleotide fasta sequences of these selected homologs are aligned using MUSCLE (Edgar RC, Nucleic Acids Research 32:1792–1797, 2004) and the alignments are visualized using Jalview (Clamp M, Cuff J, Searle SM, Barton GJ, Bioinformatics 20:426– 427, 2004).

**Fig S6** Conserved frameshift in the *fenD* gene across strains harboring group B fengycin BGCs. A. BGCs from group B of fengycins with missing domains in *fenD* gene. B. Part of the phylogenetic tree including fengycin group B strains (Complete tree in Figure S4). C. Nucleotide sequence alignment with conserved deletion at positions 3126-3127 that lead to frameshift.

**Table S1** Amino acid specificity prediction-based groups of iturin like BGCs

**Data S1** Excel table with information on all the genomes, GTDB phylogeny, MLST genes, BGCs detected and BiG-SCAPE defined GCFs along with singleton BGCs

**Data S2** Excel table with information on BGCs from families encoding fengycins, plipastatins and iturinic lipopeptides

## REFERENCES

1. Harwood CR, Mouillon J-M, Pohl S, Arnau J. 2018. Secondary metabolite production and the safety of industrially important members of the *Bacillus subtilis* group. FEMS Microbiol Rev 42:721–738.

2. Fira D, Dimkić I, Berić T, Lozo J, Stanković S. 2018. Biological control of plant pathogens by *Bacillus* species. J Biotechnol 285:44–55.

3. Kiesewalter HT, Lozano-Andrade CN, Wibowo M, Strube ML, Maróti G, Snyder D, Jørgensen TS, Larsen TO, Cooper VS, Weber T, KovácsÁ T. 2020. Genomic and chemical diversity of *Bacillus subtilis* secondary metabolites against plant pathogenic fungi. bioRxiv https://doi.org/10.1101/2020.08.05.238063.

4. Kaspar F, Neubauer P, Gimpel M. 2019. Bioactive secondary metabolites from *Bacillus subtilis*: a comprehensive review. J Nat Prod 82:2038–2053.

5. Blin K, Shaw S, Steinke K, Villebro R, Ziemert N, Lee SY, Medema MH, Weber T. 2019. antiSMASH 5.0: updates to the secondary metabolite genome mining pipeline. Nucleic Acids Res 47:W81–W87.

6. Navarro-Muñoz JC, Selem-Mojica N, Mullowney MW, Kautsar SA, Tryon JH, Parkinson EI, De Los Santos ELC, Yeong M, Cruz-Morales P, Abubucker S, Roeters A, Lokhorst W, Fernandez-Guerra A, Cappelini LTD, Goering AW, Thomson RJ, Metcalf WW, Kelleher NL, Barona-Gomez F, Medema MH. 2020. A computational framework to explore large-scale biosynthetic diversity. Nat Chem Biol 16:60–68.

7. Alanjary M, Steinke K, Ziemert N. 2019. AutoMLST: an automated web server for generating multi-locus species trees highlighting natural product potential. Nucleic Acids Res 47:W276–W282.

8. Dunlap CA, Bowman MJ, Zeigler DR. 2020. Promotion of *Bacillus subtilis* subsp. *inaquosorum, Bacillus subtilis* subsp. *spizizenii* and *Bacillus subtilis* subsp. *stercoris* to species status. Antonie Van Leeuwenhoek 113:1–12.

9. Chen XH, Koumoutsi A, Scholz R, Eisenreich A, Schneider K, Heinemeyer I, Morgenstern B, Voss B, Hess WR, Reva O, Junge H, Voigt B, Jungblut PR, Vater J, Süssmuth R, Liesegang H, Strittmatter A, Gottschalk G, Borriss R. 2007. Comparative analysis of the complete genome sequence of the plant growth-promoting bacterium *Bacillus amyloliquefaciens* FZB42. Nat Biotechnol 25:1007–1014.

10. Nishito Y, Osana Y, Hachiya T, Popendorf K, Toyoda A, Fujiyama A, Itaya M, Sakakibara Y. 2010. Whole genome assembly of a natto production strain *Bacillus subtilis* natto from very short read data. BMC Genomics 11:243.

11. Dunlap CA, Bowman MJ, Rooney AP. 2019. Iturinic lipopeptide diversity in the *Bacillus subtilis* species group - important antifungals for plant disease biocontrol applications. Front Microbiol 10:1794.

12. Wu H, Wang D, Gao F. 2020. Toward a high-quality pan-genome landscape of *Bacillus subtilis* by removal of confounding strains. Brief Bioinform bbaa013.

13. Kiesewalter HT, Lozano-Andrade CN, Maróti G, Snyder D, Cooper VS, Jørgensen TS, Weber T, Kovács Á T. 2020. Complete genome sequences of 13 *Bacillus subtilis* soil isolates for studying secondary metabolite diversity. Microbiol Resour Announc 9:e01406–19.

14. Medema MH, Kottmann R, Yilmaz P, Cummings M, Biggins JB, Blin K, de Bruijn I, Chooi YH, Claesen J, Coates RC, Cruz-Morales P, Duddela S, Düsterhus S, Edwards DJ, Fewer DP, Garg N, Geiger C, Gomez-Escribano JP, Greule A, Hadjithomas M, Haines AS, Helfrich EJN, Hillwig ML, Ishida K, Jones AC, Jones CS, Jungmann K, Kegler C, Kim HU, Kötter P, Krug D, Masschelein J, Melnik AV, Mantovani SM, Monroe EA, Moore M, Moss N, Nützmann H-W, Pan G, Pati A, Petras D, Reen FJ, Rosconi F, Rui Z, Tian Z, Tobias NJ, Tsunematsu Y, Wiemann P, Wyckoff E, Yan X, Yim G, Yu F, Xie Y, Aigle B, Apel AK, Balibar CJ, Balskus EP, Barona-Gómez F, Bechthold A, Bode HB, Borriss R, Brady SF, Brakhage AA, Caffrey P, Cheng Y-Q, Clardy J, Cox RJ, De Mot R, Donadio S, Donia MS, van der Donk WA, Dorrestein PC, Doyle S, Driessen AJM, Ehling-Schulz M, Entian K-D, Fischbach MA, Gerwick L, Gerwick WH, Gross H, Gust B, Hertweck C, Höfte M, Jensen SE, Ju J, Katz L, Kaysser L, Klassen JL, Keller NP, Kormanec J, Kuipers OP, Kuzuyama T, Kyrpides NC, Kwon H-J, Lautru S, Lavigne R, Lee CY, Linquan B, Liu X, Liu W, Luzhetskyy A, Mahmud T, Mast Y, Méndez C, Metsä-Ketelä M, Micklefield J, Mitchell DA, Moore BS, Moreira LM, Müller R, Neilan BA, Nett M, Nielsen J, O’Gara F, Oikawa H, Osbourn A, Osburne MS, Ostash B, Payne SM, Pernodet J-L, Petricek M, Piel J, Ploux O, Raaijmakers JM, Salas JA, Schmitt EK, Scott B, Seipke RF, Shen B, Sherman DH, Sivonen K, Smanski MJ, Sosio M, Stegmann E, Süssmuth RD, Tahlan K, Thomas CM, Tang Y, Truman AW, Viaud M, Walton JD, Walsh CT, Weber T, van Wezel GP, Wilkinson B, Willey JM, Wohlleben W, Wright GD, Ziemert N, Zhang C, Zotchev SB, Breitling R, Takano E, Glöckner FO. 2015. Minimum Information about a Biosynthetic Gene cluster. Nat Chem Biol 11:625–631.

15. Hagberg AA, Schult DA, Swart PJ. 2008. Exploring network structure, dynamics, and function using NetworkX, p. 11–15. In Varoquaux, G, Vaught, T, Millman, J (eds.), 7th Python in Science Conference (SciPy 2008).

16. Mohite OS, Lloyd CJ, Monk JM, Weber T, Palsson BO. Pangenome analysis of Enterobacteria reveals richness of secondary metabolite gene clusters and their associated gene sets. bioRxiv https://doi.org/10.1101/781328.

17. Katoh K, Standley DM. 2013. MAFFT multiple sequence alignment software version 7: improvements in performance and usability. Mol Biol Evol 30:772–780.

18. Capella-Gutiérrez S, Silla-Martínez JM, Gabaldón T. 2009. trimAl: a tool for automated alignment trimming in large-scale phylogenetic analyses. Bioinformatics 25:1972–1973.

19. Nguyen L-T, Schmidt HA, von Haeseler A, Minh BQ. 2015. IQ-TREE: a fast and effective stochastic algorithm for estimating maximum-likelihood phylogenies. Mol Biol Evol 32:268–274.

20. Hoang DT, Chernomor O, von Haeseler A, Minh BQ, Vinh LS. 2018. UFBoot2: Improving the ultrafast bootstrap approximation. Mol Biol Evol 35:518–522.

21. Huerta-Cepas J, Dopazo J, Gabaldón T. 2010. ETE: a python Environment for Tree Exploration. BMC Bioinformatics 11:24.

22. Parks DH, Chuvochina M, Waite DW, Rinke C, Skarshewski A, Chaumeil P-A, Hugenholtz P. 2018. A standardized bacterial taxonomy based on genome phylogeny substantially revises the tree of life. Nat Biotechnol 36:996–1004.

23. Parks DH, Chuvochina M, Chaumeil P-A, Rinke C, Mussig AJ, Hugenholtz P. 2020. A complete domain-to-species taxonomy for Bacteria and Archaea. Nat Biotechnol 38:1079–1086.

24. Virtanen P, Gommers R, Oliphant TE, Haberland M, Reddy T, Cournapeau D, Burovski E, Peterson P, Weckesser W, Bright J, van der Walt SJ, Brett M, Wilson J, Millman KJ, Mayorov N, Nelson ARJ, Jones E, Kern R, Larson E, Carey CJ, Polat İ, Feng Y, Moore EW, VanderPlas J, Laxalde D, Perktold J, Cimrman R, Henriksen I, Quintero EA, Harris CR, Archibald AM, Ribeiro AH, Pedregosa F, van Mulbregt P, SciPy 1.0 Contributors. 2020. SciPy 1.0: fundamental algorithms for scientific computing in Python. Nat Methods 17:261–272.

