## Supplementary material for "Phylogenetic distribution of secondary metabolites in the *Bacillus subtilis* species complex": Fig S1 to Fig S6 and Tables S1

SI includes:

Figures S1 to S6

Table S1

Data S1 and S2

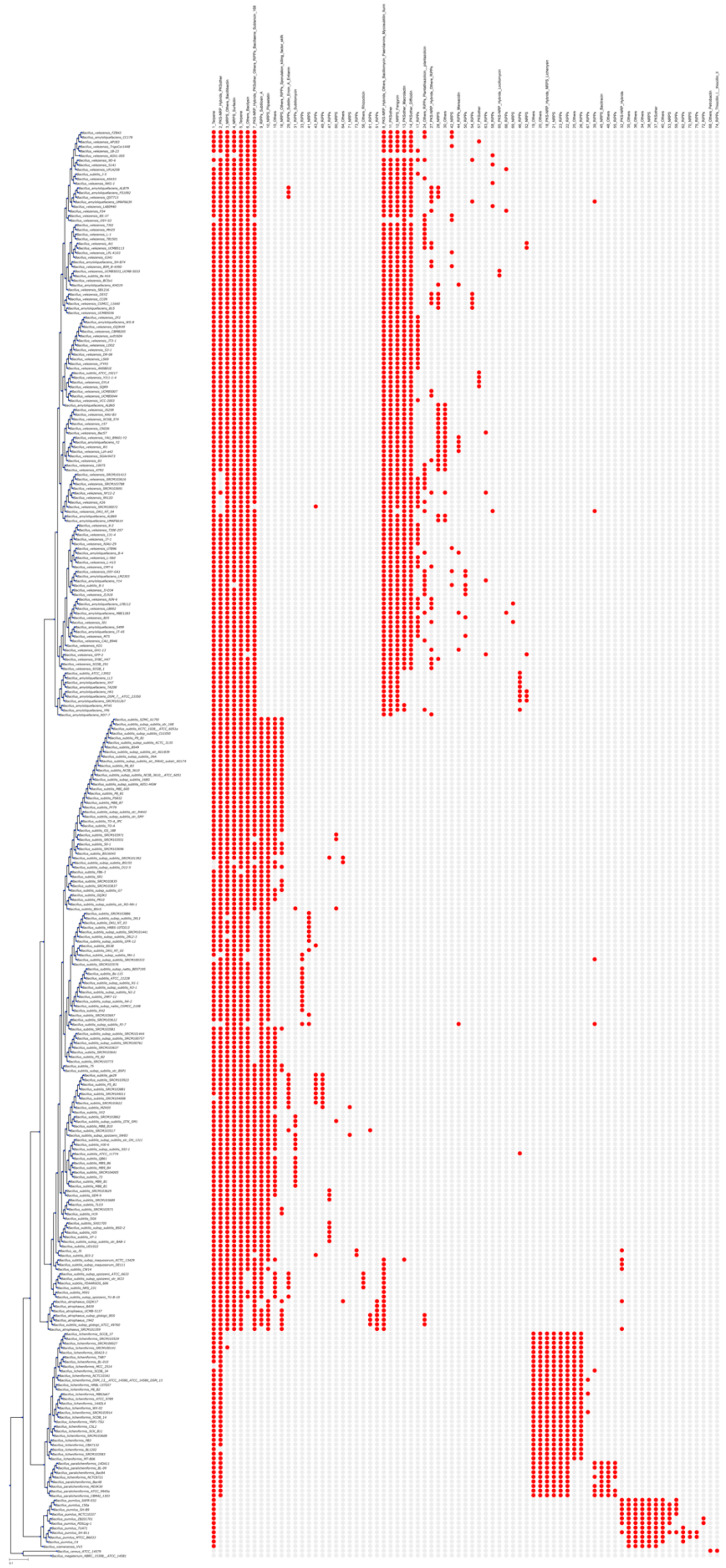

Fig. S1. Phylogenetic tree and presence absence of different BGC families across *Bacillus* group

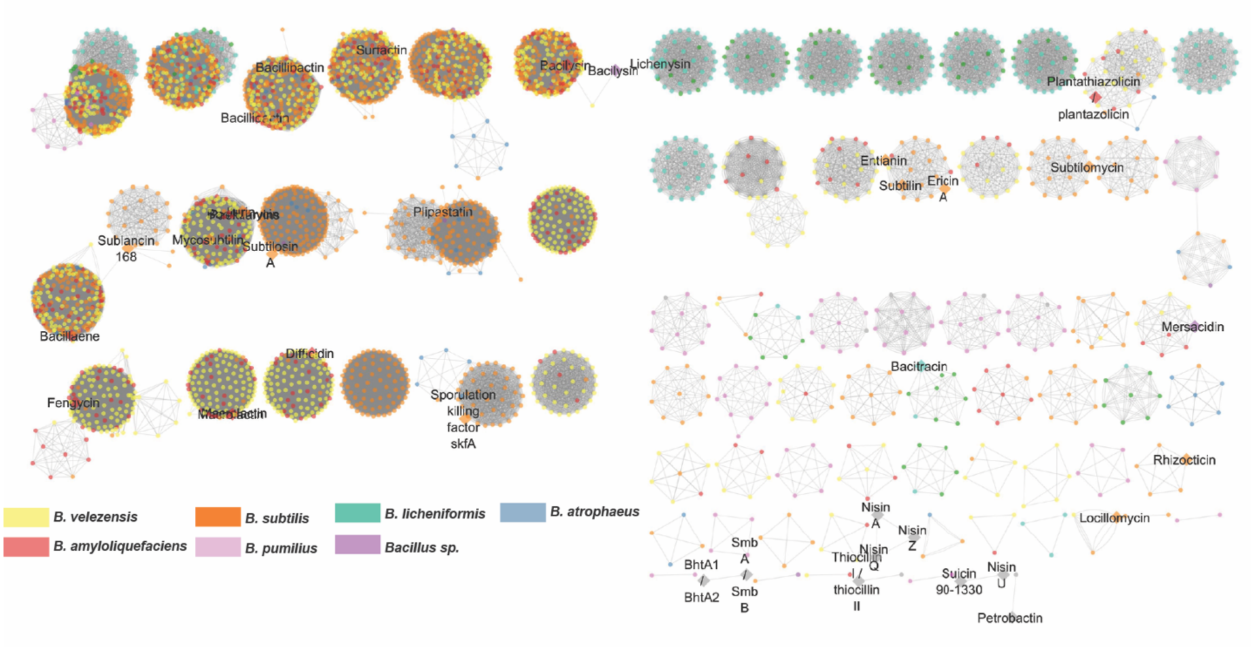


Fig. S2. Similarity network overview of all BGCs detected across *Bacillus* genome visualized using Cytoscape (Shannon P, Markiel A, Ozier O, Baliga NS, Wang JT, Ramage D, Amin N, Schwikowski B, Ideker T, Genome Res 13:2498–2504, 2003).

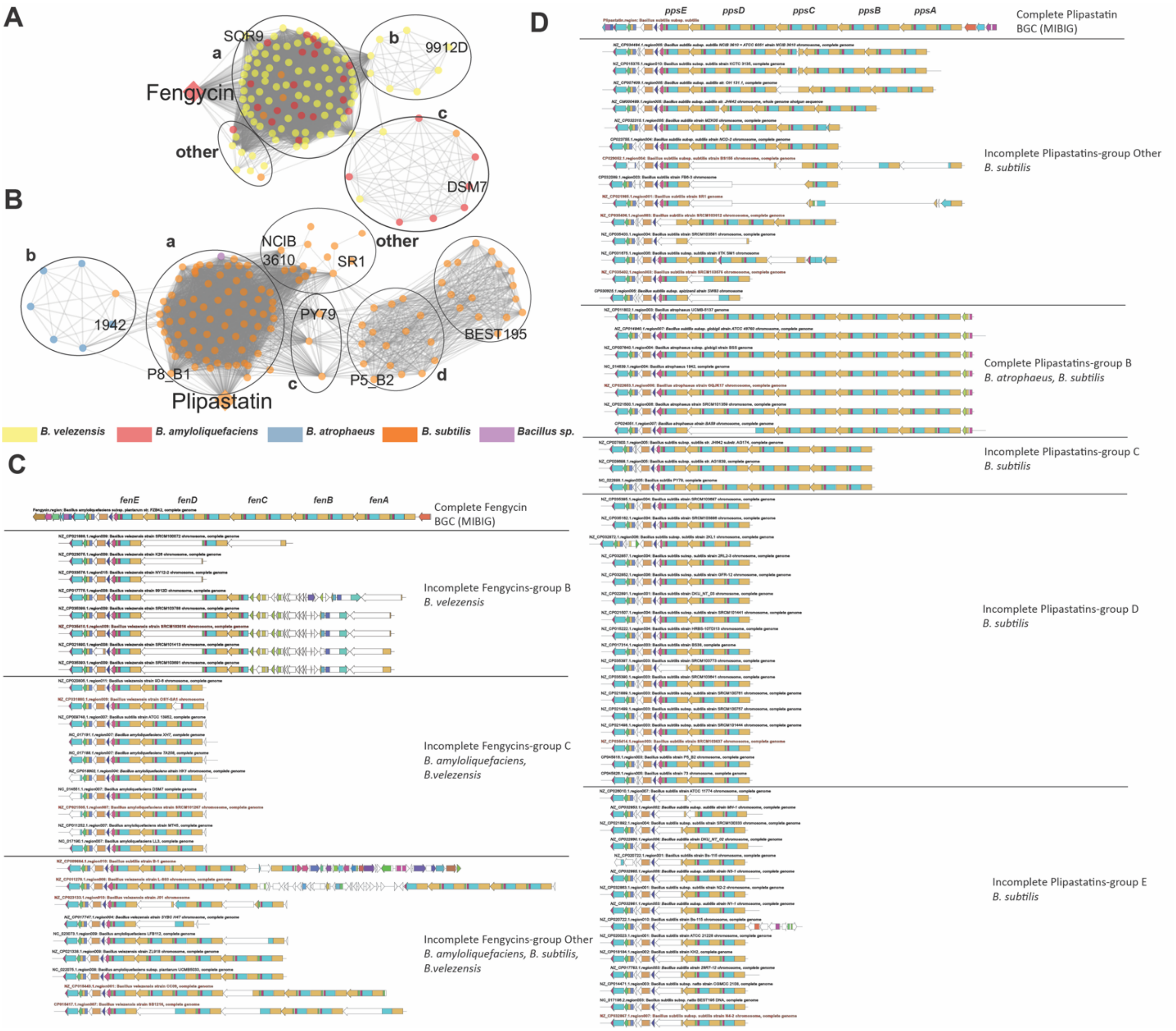


Fig. S3. Genetic structure variations in partial BGCs of plipastatin and fengycin families

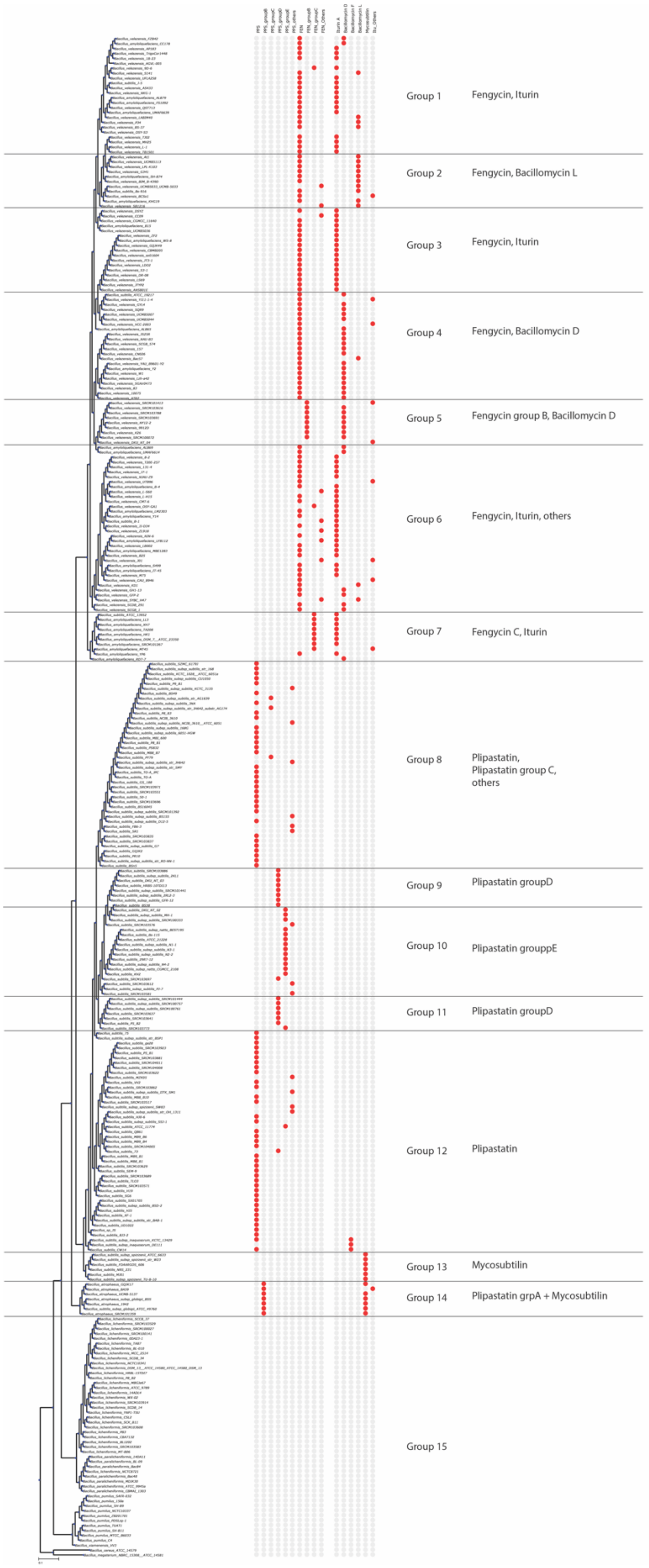


Fig. S4. Phylogenetic tree distribution of different fengycins and iturins across 310 genomes of *Bacillus sp.*

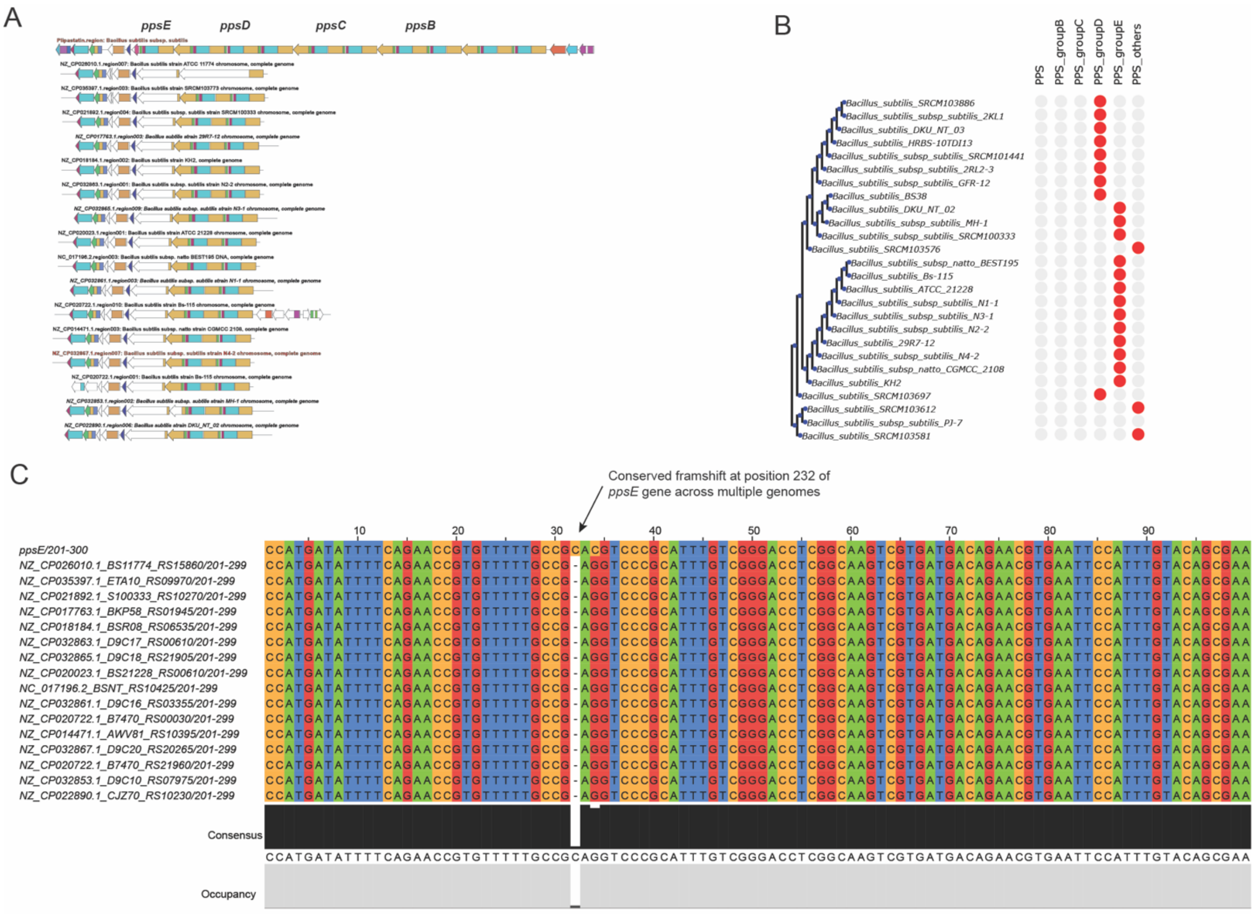


Fig. S5. Conserved frameshift in the *ppsE* gene across strains harboring group E plipastatin BGCs A. BGCs from group E of plipastatins with missing domains in *ppsE* gene. B. Part of the phylogenetic tree including plipastatin group E strains (Complete tree in Figure S4). C. Nucleotide sequence alignment with conserved deletion at position 232 that lead to frameshift. The nucleotide fasta sequences of these selected homologs are aligned using MUSCLE (Edgar RC, Nucleic Acids Research 32:1792–1797, 2004) and the alignments are visualized using Jalview (Clamp M, Cuff J, Searle SM, Barton GJ, Bioinformatics 20:426–427, 2004).

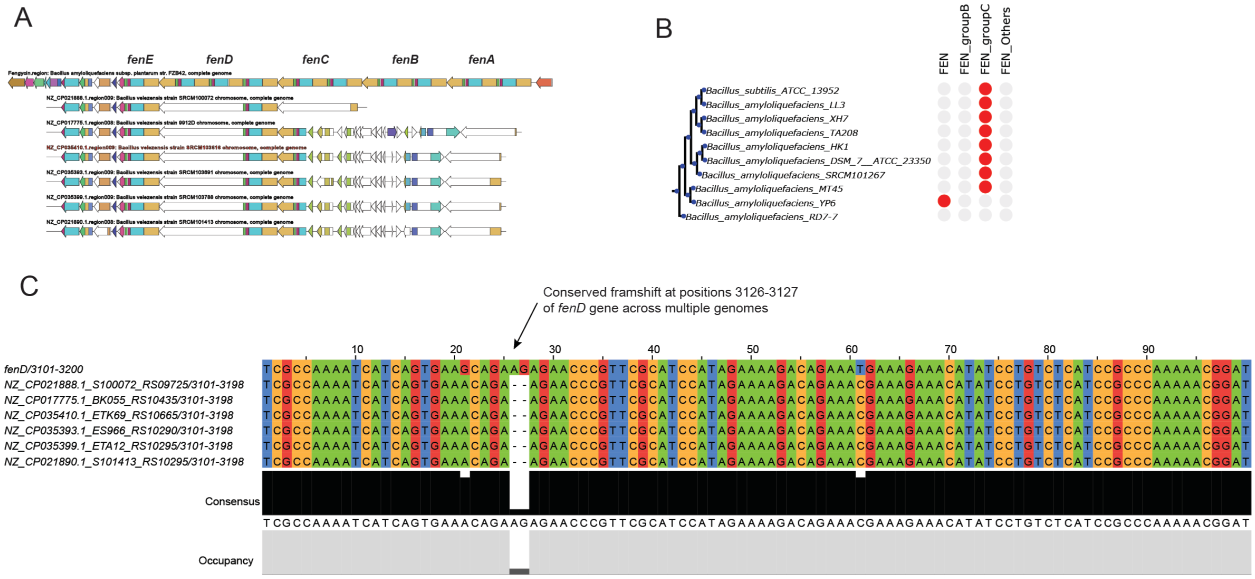


Fig. S6. Conserved frameshift in the *fenD* gene across strains harboring group B fengycin BGCs A. BGCs from group B of fengycins with missing domains in *fenD* gene. B. Part of the phylogenetic tree including fengycin group B strains (Complete tree in Figure S4). C. Nucleotide sequence alignment with conserved deletion at positions 3126-3127 that lead to frameshift.

Table S1: Amino acid specificity prediction-based groups of iturinic lipopeptide BGCs

| Group of BGC | AA prediction sequence | *B. subtilis* | *B. velezensis* | *B. amyloliquefaciens* | *B. atrophaeus* |
|---|---|---|---|---|---|
| Iturin A | ser,asn,pro,gln,asn,tyr,asn,fatty_acid,mal | 3 | 42 | 19 | 0 |
| Bacillomycin D | thr,ser,glu,pro,asn,tyr,asn,fatty_acid,mal | 1 | 28 | 6 | 0 |
| Bacillomycin F | thr,asn,pro,gln,asn,tyr,asn,fatty_acid,mal | 3 | 0 | 0 | 0 |
| Bacillomycin L | thr,ser,glu,ser,asn,tyr,asn,fatty_acid,mal | 1 | 14 | 2 | 0 |
| Mycosubtilin | asn,ser,pro,gln,asn,tyr,asn,fatty_acid,mal | 7 | 0 | 0 | 5 |
| Incomplete 1 | asn,tyr,asn,fatty_acid,mal | 0 | 2 | 0 | 0 |
| Incomplete 2 | asn,ser,tyr,asn,fatty_acid,mal,inactive | 0 | 0 | 0 | 1 |
| Incomplete 3 | ser,asn,asn,tyr,asn,fatty_acid,mal | 0 | 2 | 0 | 0 |
| Incomplete 4 | thr,ser,tyr,asn,fatty_acid,mal | 0 | 1 | 0 | 0 |
| Incomplete 5 | ser,asn,gln,asn,tyr,asn,fatty_acid,mal,inactive | 0 | 0 | 1 | 0 |
| Incomplete 6 | ser,asn,pro,gln,asn,tyr,fatty_acid,mal | 0 | 1 | 0 | 0 |
| Incomplete 7 | thr,ser,glu,pro,asn,tyr,fatty_acid,mal | 0 | 1 | 0 | 0 |
| Incomplete 8 | ser,glu,pro,asn,tyr,asn,fatty_acid,mal | 0 | 1 | 0 | 0 |

Supplementary information also includes following data:

Data S1. Excel table with information on all the genomes, GTDB phylogeny, MLST genes, BGCs detected and BiG-SCAPE defined GCFs along with singleton BGCs

Data S2. Excel table with information on BGCs from families encoding fengycins, plipastatins and iturinic lipopeptides
